# FAM134B-RHD Protein Clustering Drives Spontaneous Budding of Asymmetric Membranes

**DOI:** 10.1101/2021.01.04.425110

**Authors:** Marc Siggel, Ramachandra M. Bhaskara, Melanie K. Moesser, Ivan Đikić, Gerhard Hummer

## Abstract

Living cells constantly remodel the shape of their lipid membranes. In the endoplasmic reticulum (ER), the reticulon homology domain (RHD) of the reticulophagy regulator 1 (RETR1/FAM134B) forms dense autophagic puncta that are associated with membrane removal by ER-phagy. In molecular dynamics (MD) simulations, we find that FAM134B-RHD spontaneously forms clusters, driven in part by curvature-mediated attraction. At a critical size, the FAM134B-RHD clusters induce the formation of membrane buds. The kinetics of budding depends sensitively on protein concentration and bilayer asymmetry. Our MD simulations shed light on the role of FAM134B-RHD in ER-phagy and show that membrane asymmetry can be used to modulate the kinetics barrier for membrane remodeling.

**Graphical TOC Entry:** 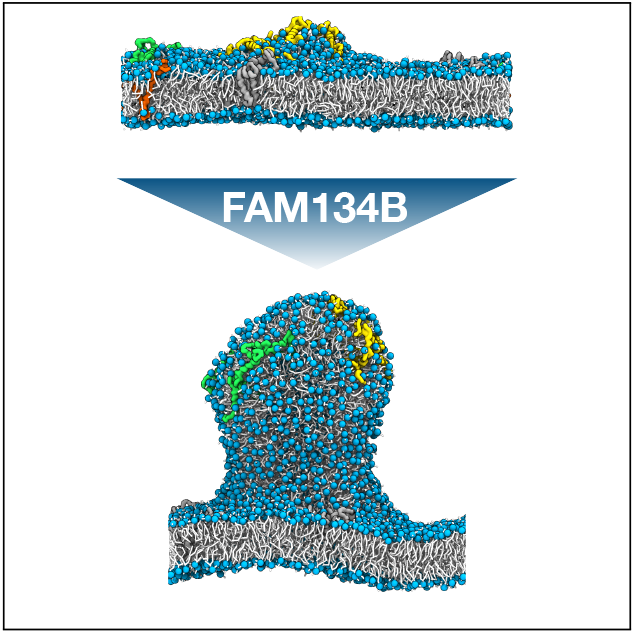

---

Cellular membranes are shaped by curvature-inducing proteins, lipid composition, or external mechanical forces.^1–3^ Protein oligomerization and clustering have been associated with membrane curvature induction, budding and scission.^4^ In the endoplasmic reticulum (ER), the reticulophagy regulator 1 (RETR1/FAM134B)^5^ localizes into the autophagic puncta associated with the control of ER size and the removal of aggregated proteins by ER-phagy.^6^ Molecular modeling and molecular dynamics (MD) simulations revealed that the reticulon homology domain (RHD) of FAM134B is responsible for membrane curvature sensing and active curvature induction.^7^ The MD simulations also showed that membrane deformation is amplified by FAM134B-RHD clustering. However, the exact mechanism and dynamics of protein clustering leading to membrane budding and scission remained unclear.

Studying the dynamics of protein-induced membrane budding has remained challenging in MD simulations. In simulations of finite membrane patches under periodic boundary conditions (PBC), large-scale fluctuations of the membrane, shape deformations, and topological changes associated with membrane scission and fusion are strongly suppressed. As alternatives, ultra coarse-grained models^8–11^ and tether pulling by external force have been used.^12^ We recently explored the use of bilayers with asymmetric leaflets for MD simulations of spontaneous membrane budding.^13^ We found that a kinetic barrier separating the metastable flat state from the stable bud shape at high leaflet asymmetry could be overcome by lateral pressure.

Here, we exploit membrane asymmetry to study membrane budding induced by clusters of FAM134B-RHD. We reasoned that wedge-shaped protein inclusions in the membrane should mimic the effects of lateral pressure. In our MD simulations (see Methods in the Supporting Information), we varied the leaflet asymmetry and protein concentration to modulate the kinetic barrier and energetic driving force for budding. We initiated simulations from flat metastable bilayers with different numbers of 1-palmitoyl-2-oleoyl-glycero-3-phosphocholine (POPC) lipids in the upper and lower leaflets, Δ*N* = *N*_upper_ – *N*_lower_ = 100, 200, 300 and 400; and different numbers of membrane-embedded FAM134B-RHDs, *n* =1 to 13. The relative asymmetries Δ*N/N* were 6.8%, 13.6%, 20.4% and 27.2%, where *N* = (*N*_upper_ + *N*_lower_)/2. For each setup, we performed three independent runs (Table S1). In all our simulations, we used the MARTINI^14^ coarse-grained model.

We found that FAM134B-RHD proteins are capable of reshaping a membrane from a flat to a budded shape (Figure 1, Supporting Movie S1). Budding is associated with a sharp decrease in the box width *L_x_* as membrane area is absorbed into the nascent bud (Figure 1A), making *L_x_* an excellent reporter on membrane shape changes. Figure 1B shows the initial flat membrane and final budded state in an MD simulation with *n* = 9 FAM134B-RHDs and a leaflet asymmetry of Δ*N* = 300. We had shown earlier that without proteins, the POPC membrane remained trapped in the flat metastable state for over 7 *μ*s in a system with Δ*N*/*N* = 0.304 and *N* = 1634 (Figure S1A in ref 13), i.e., at an asymmetry higher than any of the systems studied here.

**Figure 1:**
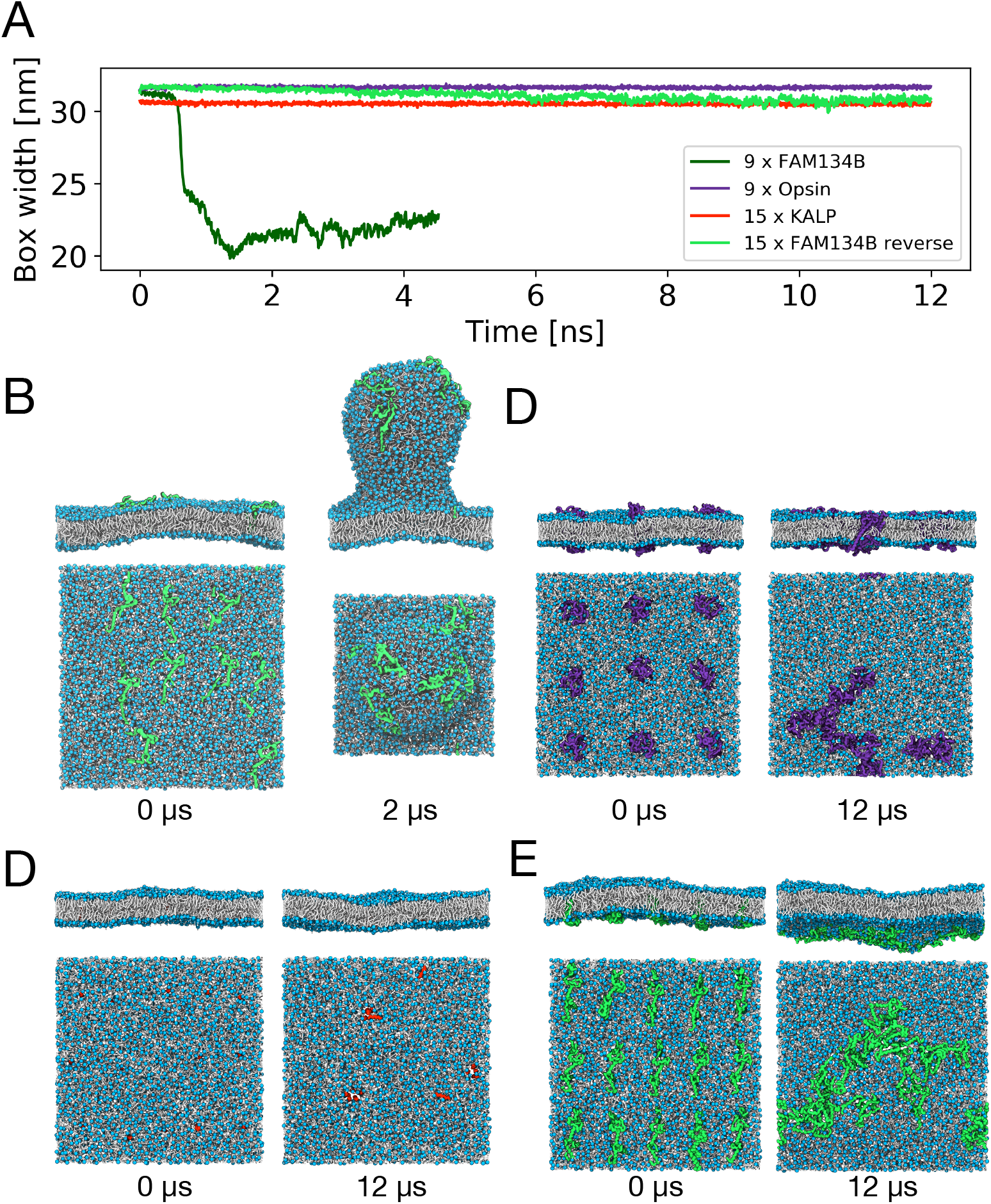
FAM134B-RHD specifically induces membrane budding in asymmetric membranes. (A) Time traces of box width for one exemplary budding event with nine FAM134B-RHD (blue curve) and three control simulations with nine opsins (purple), 15 KALP_15_ (red), and 15 FAM134B-RHD in reverse topology (green). Corresponding snapshots (top and side views) at times 0 and 12 *μ*s are shown in B-E. All systems had an asymmetry of Δ*N* = 300. (B) Nine FAM134 proteins embedded in asymmetric bilayer induce the formation of a membrane bud within 2 *μ*s. Simulation snapshots of (C) nine opsin proteins, (D) 15 KALP_15_ peptides, and (E) 15 FAM134B-RHD (inverted topology), which do not induce budding of the asymmetric membrane. Embedded proteins are shown in surface representation, lipid phosphate groups in blue, and lipid tails in white. Water and ions are omitted for clarity.

We confirmed that membrane reshaping and budding are specific to the curvature-inducing properties of FAM134B-RHD and not triggered by membrane inclusions alone (Figure 1). As a control, we performed simulations of proteins not associated with membrane remodeling: (i) the transmembrane helical peptide KALP_15_, whose hydrophobic mismatch resembles that of the transmembrane hairpins of FAM134B-RHD; and (ii) the 7-transmembrane helical G-protein coupled receptor opsin. In these control simulations, the bilayers remained nearly flat (Figure 1C, D). As an additional control, we placed *n* =15 FAM134B-RHD proteins into the membrane in reverse topology, i.e., with the N and C termini on the lower side of the membrane. In effect, this inverts the sign of the membrane asymmetry to Δ*N* = −300. Even *n* =15 FAM134B-RHD proteins were not sufficient to induce budding at this unfavorable asymmetry (Figure 1E). The protein identity and the insertion topology are thus decisive factors for budding.

Higher asymmetry and protein concentration both accelerate the induction of membrane buds (Figure 2). In simulations with low leaflet asymmetry, Δ*N* = 100, we did not observe any major membrane shape changes (Figure 2) at any protein concentration. At this low value of leaflet asymmetry, the excess lipids can be accommodated by minor compression and expansion of the respective leaflets of a flat bilayer. As the FAM134B-RHD proteins diffused in the membrane plane and formed transient clusters, the associated membrane deformations remained local and did not induce any budding events (Figure 3, Δ*N* = 100). At intermediate leaflet asymmetry (Δ*N* = 200, 300) and low FAM134B-RHD concentration (n ≤ 5), we again did not observe budding events (Figure 2). However, by increasing the number of FAM134B-RHD proteins (*n* > 5), we observed spontaneous membrane budding, indicating that a critical number of proteins is required (Figure 3; Δ*N* = 200, 300). At high leaflet asymmetry (Δ*N* = 400), we observed the largest number of spontaneous budding transitions from the flat bilayer, indicating that the kinetic barrier was lowered significantly even at low protein concentrations (Figure 2). For Δ*N* = 400, we observed a budding event already with a single FAM134B-RHD protein embedded in the bilayer.

**Figure 2:**
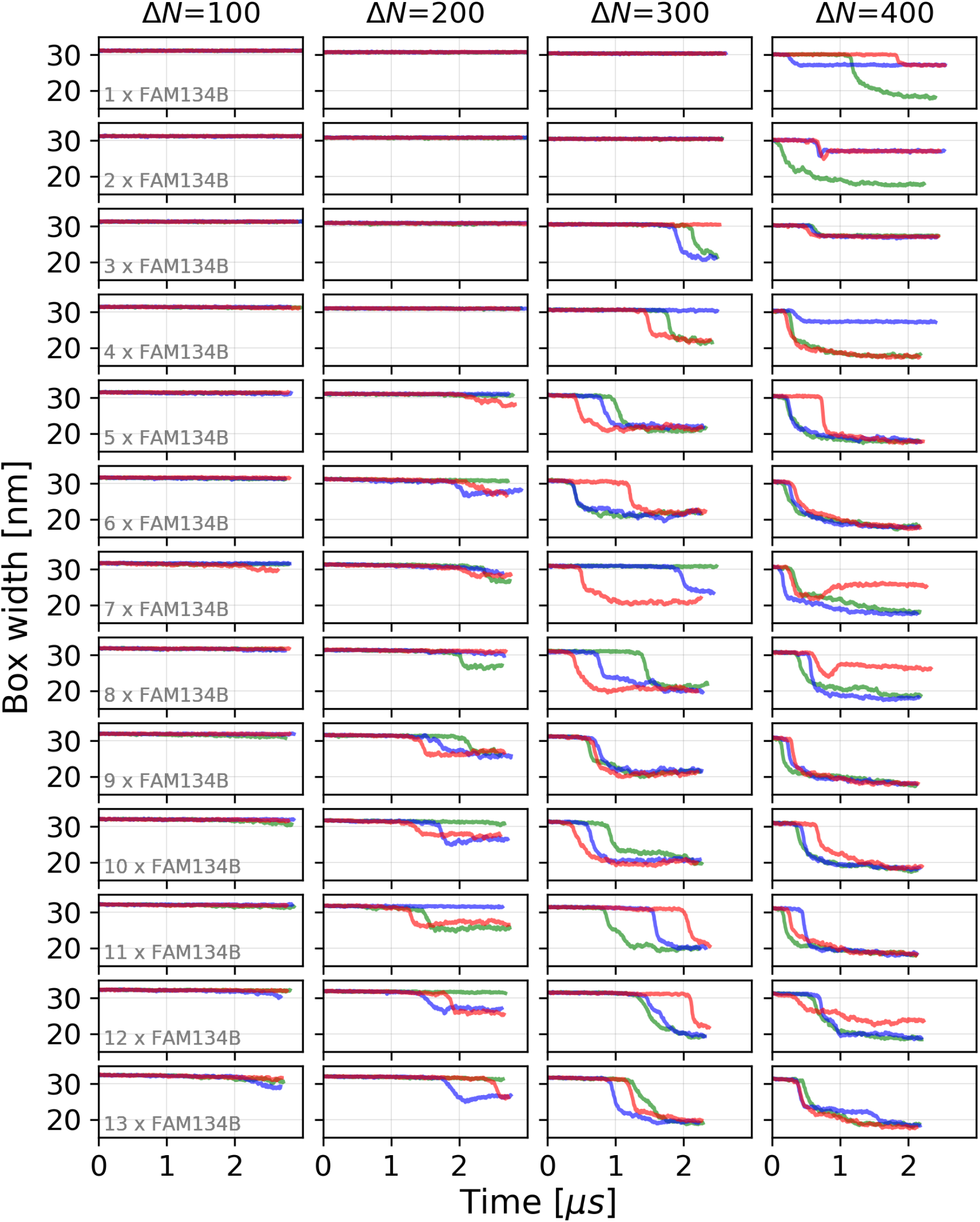
Box width as the reporter of membrane shape changes. Time traces of the box width *L_x_* are shown for MD simulations of *n* = 1-13 FAM134B-RHD proteins (top to bottom) in POPC membranes with leaflet asymmetries of Δ*N* = 100, 200, 300 and 400 (left to right). The three replicates are distinguished by color.

**Figure 3:**
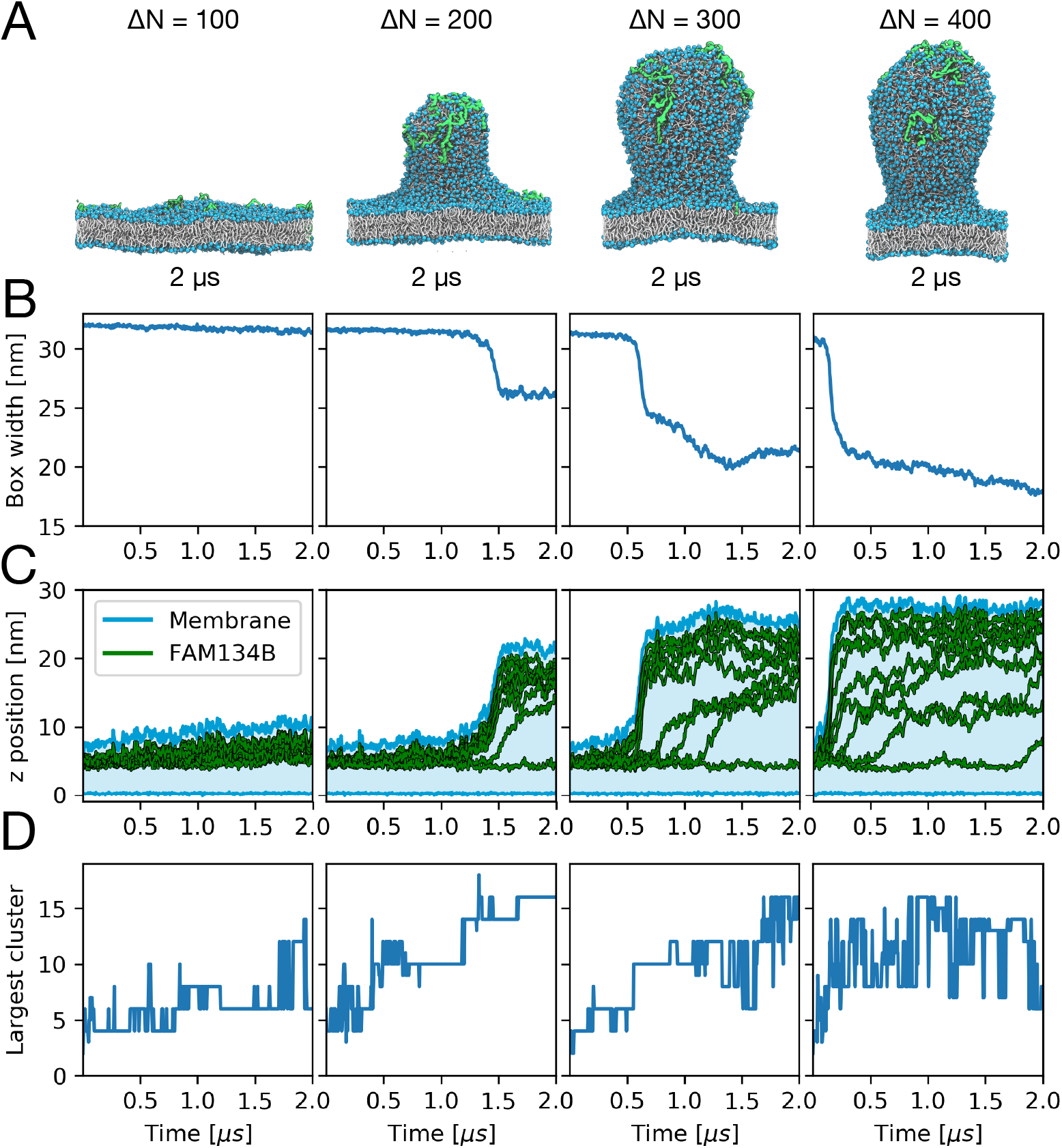
Budding of membranes with different leaflet asymmetries. Results are shown for asymmetries of Δ*N* = 100, 200, 300 and 400 (left to right) with *n* = 9 FAM134B-RHD proteins in the membrane. (A) Snapshots of asymmetric membranes after 2 *μ*s with FAM134B-RHD shown in green, lipid headgroups in blue, and lipid tails in white. Water and ions are omitted for clarity. (B) Time traces of box widths *L_x_*. (C) Vertical displacement z of individual FAM134B-RHD proteins (center-of-mass positions; green lines). The highest and lowest points in the membrane are shown as blue lines and the intervening range in light blue shading. (D) Size of the largest FAM134B-RHD cluster as a function of time for different membrane asymmetries. Transmembrane helical hairpins were clustered and counted individually.

Along with the increase in the number of budding events, the waiting times for budding decreased by increasing FAM134B-RHD concentration (Figure 2). Importantly, though, in all cases there was a time lag between the start of the simulations and budding, indicating that even at the highest asymmetry and largest protein concentrations, budding had to overcome a kinetic barrier.

The shape of the buds depends on membrane asymmetry. The tubular shape of the buds formed at lower asymmetry (Δ*N* = 200) changed into a more spherical shape at higher asymmetry (Δ*N* = 300, 400; Figure 3A). This shape change is reflected in the differences in box width after budding (Figure 3B). With increasing asymmetry, the box length decreases more strongly, indicating that a larger membrane area is absorbed into the bud. At the highest asymmetry (Δ*N* = 400), the bud size is likely limited by self-interactions of the nascent bud across the periodic boundaries of the simulation box.

At high leaflet asymmetry (Δ*N* = 400), we observed an alternate route to alleviate membrane stress. In a few replicates, mostly at low protein concentrations, the excess lipids of the dense upper leaflet folded onto themselves to create a bicelle-like protrusion attached to the otherwise flat bilayer (Figure 4C). The resulting drop in the box width was less pronounced than with actual budding (Figures 2 and 4A). FAM134B-RHD proteins localized to the connection between the protrusion and the membrane, but did not sort onto the protrusion, unlike the vesicle buds.

**Figure 4:**
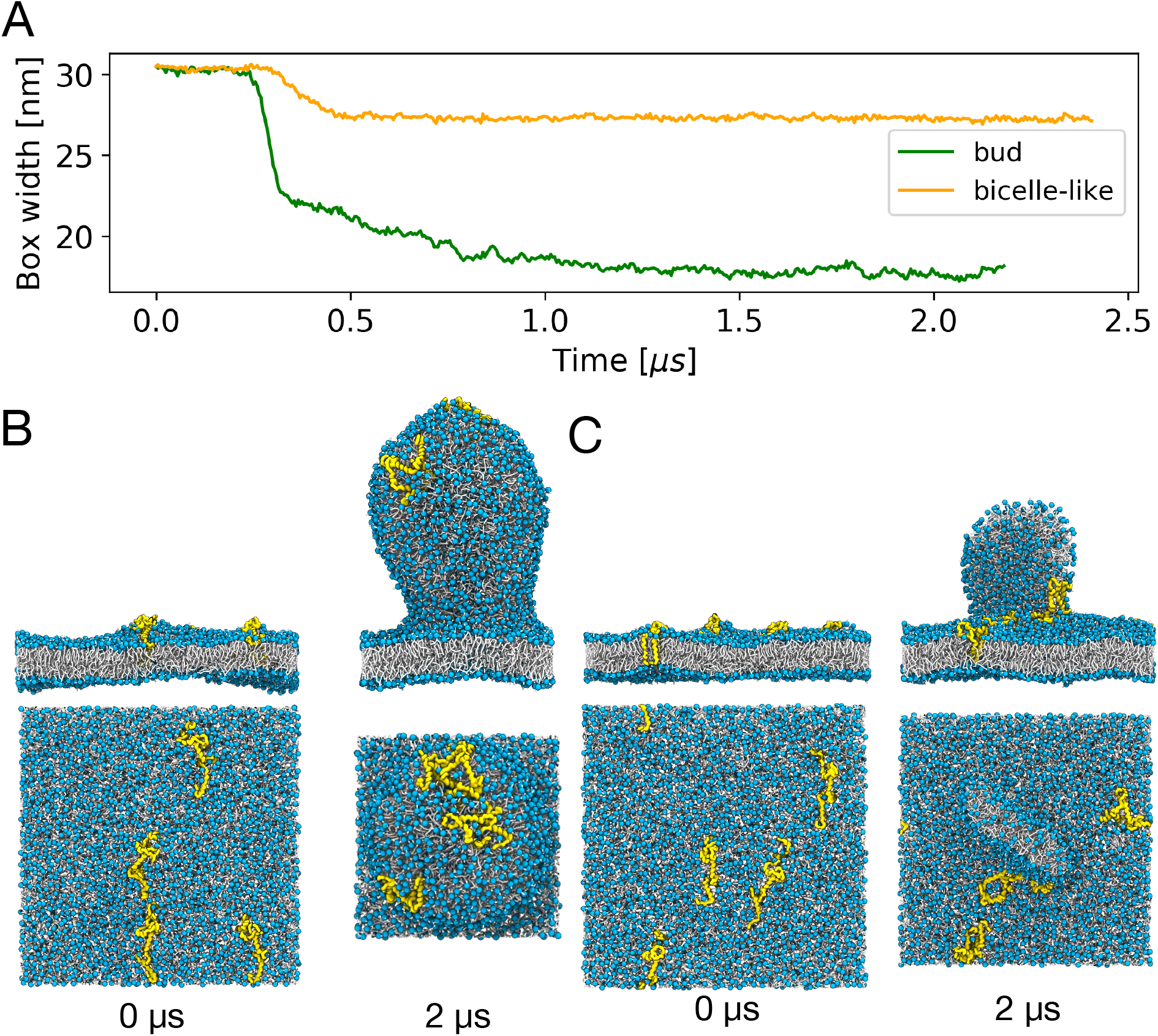
Different pathways to alleviate asymmetric membrane stress. (A) Time traces of the box width *L_x_* in two MD simulations with *n* = 4 FAM134B-RHD proteins and an asymmetry of Δ*N* = 400 that led to the formation of a membrane bud (green) and a bicelle-like protrusion (orange), respectively. (B,C) Beginning (0 *μ*s; left) and end states (2 *μ*s; right) with (B) a membrane bud and (C) a bicelle-like protrusion, respectively. FAM134B-RHD proteins are shown in yellow, and POPC lipids with blue headgroups and white acyl chains.

FAM134B-RHD clusters act cooperatively to induce budding, as illustrated in Figure 3 for systems with *n* = 9 FAM134B-RHD proteins and different leaflet asymmetries Δ*N*. By monitoring the vertical displacement *z* of the centers of mass of individual proteins with respect to the lowest lipid headgroup in the lower leaflet (Figure 3C), we found that seven, five, and three of the nine proteins were directly involved in the budding events at Δ*N* = 200, 300 and 400, respectively. The sharp increase in the z positions of these proteins correlates with the increase in membrane height and the contraction of the box (Figure 3C). We then noticed that these proteins formed a distinct cluster, whose size had increased until just before budding (Figure 3D). Small lipid number asymmetries, Δ*N* = 200, required a larger cluster of about 15 FAM134B-RHD transmembrane hairpins than larger asymmetries, Δ*N* = 300, where about ten hairpins sufficed. For Δ*N* = 400, already a single FAM134B-RHD could induce budding. The observations that cluster formation preceded budding and that the required cluster size increases with decreasing energetic driving force together indicate that FAM134B-RHD cluster formation is critical for budding.

The critical role of FAM134B-RHD clusters is confirmed by their localization on the emerging bud. As illustrated in a detailed view of the time evolution of FAM134B-RHD clusters during a representative budding event for Δ*N* = 300 and *n* = 9 (Figure 5), we found that budding is initiated at the site of a pre-formed FAM134B-RHD cluster (yellow, consisting of three proteins). Other FAM134B-RHD clusters (with two proteins each) merged with this large cluster (Figure 5A, B), which then triggered the rapid emergence of a membrane bud. The FAM134B-RHD clusters occupied the cusp of the emerging bud and remained there during the entire transition, indicating that the active curvature induction of the emergent clusters lowers the kinetic barrier for budding (Figure 5B).

**Figure 5:**
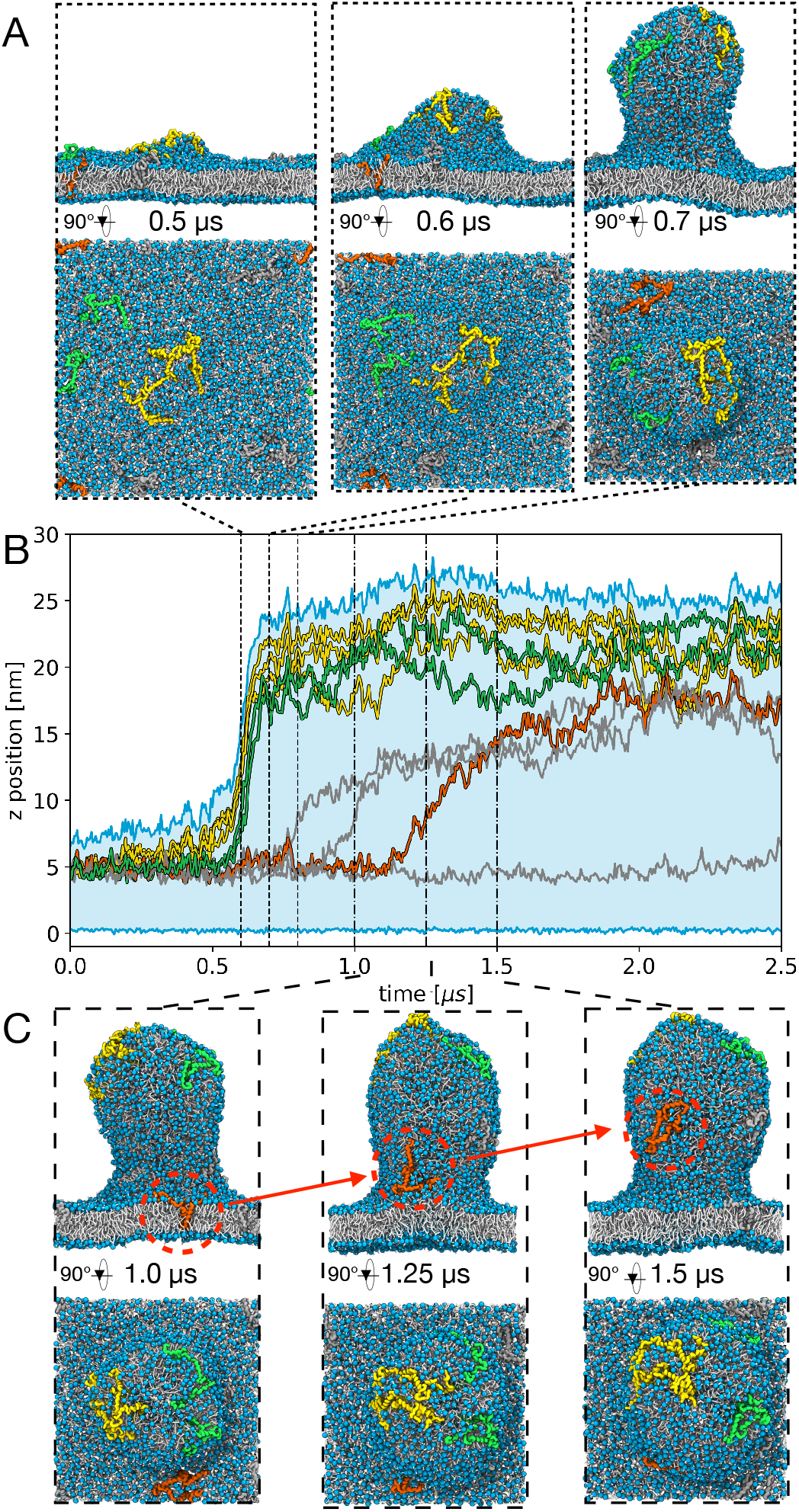
Detailed view of FAM134B-RHD clustering, membrane budding and curvature segregation. Results are shown for *n* = 9 FAM134B-RHD proteins and an asymmetry of Δ*N* = 300, corresponding to Supporting Movie S1. (A) Simulation snapshots at critical time points during the spontaneous formation of a membrane bud. (B) Vertical displacement *z* of the embedded proteins (center-of-mass positions: yellow, green, red) and the emerging bud (highest and lowest lipid headgroups of the two leaflets: blue lines; light blue shading in between) as function of time. Yellow and green traces indicate clusters of FAM134B-RHD involved in bud formation. (C) Simulation snapshots separated by 250 ns highlight curvature-mediated segregation of a single FAM134B-RHD protein (orange; see also B) onto the pre-existing bud.

FAM134B-RHD clustering is enhanced by curvature sensing. By tracking the cluster centers with respect to time, we found that FAM134B-RHD alone or in clusters of up to three proteins preferentially diffused towards the emerging buds. Even after the emergence of the bud, the remaining FAM134B-RHD clusters and individual proteins (orange, Figure 5B, C) tend to sort onto the highly curved membrane bud. This curvature sorting is observed in all analyzed systems (Figure 3C). As time progresses, all proteins eventually migrate onto the bud and remain there. Effective attractions between membrane-embedded proteins mediated by elastic deformations of the bilayer have been studied by continuum theoretical models, molecular simulations and experiments. Protein induced curvature fields can exert long-range attractive forces to enable self-organization into clusters.^9,15^

The curvature-mediated clustering of FAM134B-RHDs is consistent with its segregation into the curved regions of the ER.^5,7^ Recent experiments indicate that FAM134B-RHD forms higher-order oligomers during ER-phagy, ^16^ in the range of cluster sizes seen here to drive membrane remodeling.

Sensors of curvature, as well as mechanical drivers of budding, are the wedge-shaped transmembrane hairpins and their flanking amphipathic helices embedded in the upper leaflet. ^7^ As a consequence of this asymmetric shape, a membrane-inserted FAM134B-RHD protein displaces about 20 POPC lipids more from the upper leaflet than from the lower leaflet (Figure S1). The wedging associated with this asymmetric footprint appears to amplify the stress caused by membrane asymmetry in our simulations. As we could show, by combining these two effects and by concentrating them locally in the membrane at a FAM134B-RHD cluster, the kinetic barrier to budding can be overcome on an MD time scale. In particular, transmembrane hairpin clustering precedes the budding transitions (Figure 3D). In vitro deletion and liposome remodeling experiments confirmed the importance of inter-hairpin interactions in curvature induction and sensing. ^7^

The curvature-sensing and curvature-inducing properties of FAM134B-RHD are instrumental to the organization of autophagic receptors in the peripheral ER. We found (i) that FAM134B-RHD proteins tend to cluster in highly curved regions of the membrane and (ii) that these clusters aid in the spontaneous formation and stabilization of large-scale membrane buds. Both observations are relevant to the biology of FAM134B-RHD in ER-phagy. First, our observation of curvature-mediated FAM134B-RHD clustering is consistent with its segregation into the curved regions of the ER.^5,7^ The formation of FAM134B-RHD clusters provides a physicochemical mechanism for the localization of FAM134B-RHD in autophagic puncta during ER-phagy, as seen in immunofluorescence experiments.^5^ The reported oligomerization of FAM134B-RHD in ER-phagy^16^ are in the range of cluster sizes seen here to drive membrane remodeling. Second, our observation of FAM134B-RHD-induced membrane budding explains the “membrane shredding” action of FAM134B-RHD, which results in the formation of tiny vesicular structures. ^5,7^ Highlighting the biological relevance of FAM134B-RHD in ER-phagy, we note that Zika and Dengue viruses proteolytically target FAM134B-RHD to evade host responses during infection.^17^

Based on our findings for FAM134B-RHD, we expect MD simulations with asymmetric membranes to be useful to screen also other proteins for possible curvature induction and sensing properties. The power to modulate the energetic driving force and the kinetic barrier for membrane shape changes appears to be exploited also in living cells, where asymmetry-creating lipid flippases have been implicated in budding processes.^18^ The observed interplay between membrane asymmetry and curvature-inducing proteins has important biological implications on how cells can regulate and induce the formation of membrane buds.

## Acknowledgement

This research was supported by the Max Planck Society (M.S., M. M., R.M.B., and G.H.), the Landes-Offensive zur Entwicklung Wissenschaftlich-ökonomischer Exzellenz (LOEWE) DynaMem program of the state of Hesse (M.S. and G.H.) and by the Deutsche Forschungsgemeinschaft (DFG, German Research Foundation) – Project number 259130777 – SFB 1177 (R.M.B, G.H. and I.D.).

## Supporting Information Available

Supporting file (PDF) contains extended methods, Table S1, Figure S1, and supporting movie legend. Supporting Movie 1: MD simulation trajectory of FAM134B-induced membrane budding event. (MP4).

