## Supporting Information for "FAM134B-RHD Protein Clustering Drives Spontaneous Budding of Asymmetric Membranes"

---

This document contains

- Extended Methods
- Supporting Table and Figure
- Supporting Movie Legend

---

### Extended Methods

**System setup and MD Simulations.** We used the MARTINI[1] coarse-grained model for the MD simulations. Coarse-grained 1-palmitoyl-2-oleoyl-glycero-3-phosphocholine (POPC) bilayers with varying degrees of asymmetry ($\Delta N = 100, 200, 300, 400$) were first built using the *insane.py* script.[2] The bilayers were then solvated with coarse-grained water containing 150 mM NaCl. All protein structures (FAM134B, $KALP_{15}$, opsins) were coarse-grained using the *martinize.py* script. For FAM134B, we used the coarse-grained model with secondary structure restraints as previously reported.[3] Control simulations were performed with $KALP_{15}$ and opsin. The $KALP_{15}$ peptide was modeled as an alpha helix with secondary structure restraints. The coarse-grained opsin molecules were modeled based on a crystal structure (PDB ID: 4J4Q) and used an elastic network with the default cutoff of 0.9 nm[4] to maintain the tertiary structure. All proteins were embedded in the asymmetric membranes ($30 \times 30 \times 18$ nm$^3$) in a square grid and energy minimized using a soft-core potential and steepest-decent algorithm to remove steric-clashes with lipids.

MD simulations were then performed using GROMACS (v. 2018.7).[5] We used the MARTINI force field (v.2.2.),[1,6] a 20 fs time-step, and a pair-interaction cutoff of 1.1 nm. Electrostatic interactions were modeled using the Coulomb potential with a reaction field. A dielectric constant of 15 was used to model interactions beyond the cutoff distance $r_c = 1.1$ nm. Electrostatic interactions were shifted to zero between 0 and 1.1 nm. Short-range Lennard-Jones interactions were also cut off at a distance of 1.1 nm and shifted to zero between 0.9 to 1.1 nm. The default Verlet-buffer cutoff scheme was used. The system temperature and pressure were maintained at 1 bar and 310 K, respectively, by using the velocity-rescale thermostat[7] and the Parrinello-Rahman barostat.[8] The Berendsen barostat was used for equilibration.[9] Production runs were simulated for at least 2 $\mu s$ for data collection and analysis in all cases. Table S1 lists all systems built and simulated.

**Analysis of MD trajectories.** The simulation trajectories and individual frames were visualized using VMD.[10] Analysis scripts using MDAnalysis (v0.19)[11,12] were implemented in python 3.7 and plots were generated using matplotlib (v.3.1),[13] SciPy (v1.2), and NumPy (v1.16).[14,15] Lipids corresponding to individual leaflets were analyzed using the LeafletFinder algorithm in MDAnalysis.[11] To analyze the segregation of the proteins into the emerging membrane bud, the entire system was translated such that the lowest point of the lower leaflet ($N_{\text{upper}} > N_{\text{lower}}$) was set to $z = 0$ nm and centered in the $xy$ plane using the highest point of the membrane for each frame. By tracing the highest and lowest $z$-positions of the phosphate (PO4) beads, we were able to locate and measure the extent of budding from the flat part of the bilayer. We monitored the time-traces of the $z$-component of the protein centers of mass to ascertain their involvement in (i) curvature induction, (ii) cluster formation, (iii) nascent stages of budding and (iv) segregation to the bud.

**Analysis of FAM134B clusters.** We quantified the FAM134B interactions by clustering of the transmembrane helical hairpins described in ref 3. Each hairpin was treated as an individual entity to account for the dynamic nature of the inter-hairpin connection.[3] Two hairpins were considered to be part of the same cluster if their centers of mass were within a distance of 3.5 nm.

Table S1: MD simulations. Listed are the number of proteins $n$, the lipid number asymmetry $\Delta N$, the number of lipids in the lower ($N_{\text{lower}}$) and upper leaflets ($N_{\text{upper}}$), the system dimensions $L_x \times L_y \times L_z$ at the start of the simulation, the simulation time, the number of replicate simulations of each setup, and the number of observed budding events (with bicelle formation in parentheses). The bottom block lists the simulations with other proteins and with FAM134B in reverse topology.

| $n$ | $\Delta N$ ($N_{\text{lower}}/N_{\text{upper}}$) | System size [nm$^3$] | Simulation time [$\mu$s] | Replicates | budding (bicelle) |
|---|---|---|---|---|---|
| 1 | 100 (1421/1521) | 30×30×18 | >2 | 3 | 0 |
| 2 | 100 (1421/1521) | 30×30×18 | >2 | 3 | 0 |
| 3 | 100 (1421/1521) | 30×30×18 | >2 | 3 | 0 |
| 4 | 100 (1421/1521) | 30×30×18 | >2 | 3 | 0 |
| 5 | 100 (1421/1521) | 30×30×18 | >2 | 3 | 0 |
| 6 | 100 (1421/1521) | 30×30×18 | >2 | 3 | 0 |
| 7 | 100 (1421/1521) | 30×30×18 | >2 | 3 | 0 |
| 8 | 100 (1421/1521) | 30×30×18 | >2 | 3 | 0 |
| 9 | 100 (1421/1521) | 30×30×18 | >2 | 3 | 0 |
| 10 | 100 (1421/1521) | 30×30×18 | >2 | 3 | 0 |
| 11 | 100 (1421/1521) | 30×30×18 | >2 | 3 | 0 |
| 12 | 100 (1421/1521) | 30×30×18 | >2 | 3 | 0 |
| 13 | 100 (1421/1521) | 30×30×18 | >2 | 3 | 0 |
| 1 | 200 (1321/1521) | 30×30×18 | >2 | 3 | 0 |
| 2 | 200 (1321/1521) | 30×30×18 | >2 | 3 | 0 |
| 3 | 200 (1321/1521) | 30×30×18 | >2 | 3 | 0 |
| 4 | 200 (1321/1521) | 30×30×18 | >2 | 3 | 0 |
| 5 | 200 (1321/1521) | 30×30×18 | >2 | 3 | 0 |
| 6 | 200 (1321/1521) | 30×30×18 | >2 | 3 | 0 |
| 7 | 200 (1321/1521) | 30×30×18 | >2 | 3 | 0 |
| 8 | 200 (1321/1521) | 30×30×18 | >2 | 3 | 1 |
| 9 | 200 (1321/1521) | 30×30×18 | >2 | 3 | 3 |
| 10 | 200 (1321/1521) | 30×30×18 | >2 | 3 | 2 |
| 11 | 200 (1321/1521) | 30×30×18 | >2 | 3 | 2 |
| 12 | 200 (1321/1521) | 30×30×18 | >2 | 3 | 2 |
| 13 | 200 (1321/1521) | 30×30×18 | >2 | 3 | 2 |
| 1 | 300 (1221/1521) | 30×30×18 | >2 | 3 | 0 |
| 2 | 300 (1221/1521) | 30×30×18 | >2 | 3 | 0 |
| 3 | 300 (1221/1521) | 30×30×18 | >2 | 3 | 2 |
| 4 | 300 (1221/1521) | 30×30×18 | >2 | 3 | 2 |
| 5 | 300 (1221/1521) | 30×30×18 | >2 | 3 | 3 |
| 6 | 300 (1221/1521) | 30×30×18 | >2 | 3 | 3 |
| 7 | 300 (1221/1521) | 30×30×18 | >2 | 3 | 2 |
| 8 | 300 (1221/1521) | 30×30×18 | >2 | 3 | 3 |
| 9 | 300 (1221/1521) | 30×30×18 | >2 | 3 | 3 |
| 10 | 300 (1221/1521) | 30×30×18 | >2 | 3 | 3 |
| 11 | 300 (1221/1521) | 30×30×18 | >2 | 3 | 3 |
| 12 | 300 (1221/1521) | 30×30×18 | >2 | 3 | 3 |
| 13 | 300 (1221/1521) | 30×30×18 | >2 | 3 | 3 |
| 1 | 400 (1121/1521) | 30×30×18 | >2 | 3 | 1(2) |
| 2 | 400 (1121/1521) | 30×30×18 | >2 | 3 | 1(2) |
| 3 | 400 (1121/1521) | 30×30×18 | >2 | 3 | 3(3) |
| 4 | 400 (1121/1521) | 30×30×18 | >2 | 3 | 2(1) |
| 5 | 400 (1121/1521) | 30×30×18 | >2 | 3 | 3 |
| 6 | 400 (1121/1521) | 30×30×18 | >2 | 3 | 3 |
| 7 | 400 (1121/1521) | 30×30×18 | >2 | 3 | 2(1) |
| 8 | 400 (1121/1521) | 30×30×18 | >2 | 3 | 2(1) |
| 9 | 400 (1121/1521) | 30×30×18 | >2 | 3 | 3 |
| 10 | 400 (1121/1521) | 30×30×18 | >2 | 3 | 3 |
| 11 | 400 (1121/1521) | 30×30×18 | >2 | 3 | 3 |
| 12 | 400 (1121/1521) | 30×30×18 | >2 | 3 | 3 |
| 13 | 400 (1121/1521) | 30×30×18 | >2 | 3 | 3 |
| 9(opsins) | 300 (1221/1521) | 30×30×18 | 12 | 1 | 0 |
| 15(KALP$_{15}$ | 300 (1221/1521) | 30×30×18 | 12 | 1 | 0 |
| 15(reverse) | 300 (1221/1521) | 30×30×18 | 12 | 1 | 0 |

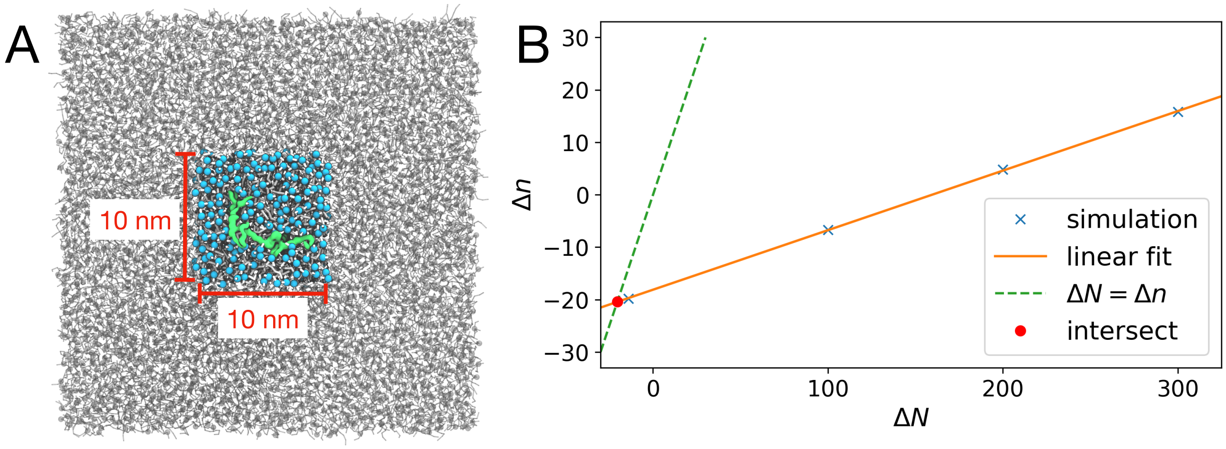


Figure S1: Asymmetric footprint of FAM134B in the membrane. (A) Top view of a system with a single FAM134B-RHD molecule in an asymmetric membrane ($\Delta N = 100$). The blue lipids indicate the $10 \times 10$ nm$^2$ square around the center of mass of FAM134B-RHD (green), within which lipid numbers $n_{\text{upper}}$ and $n_{\text{lower}}$ in the two leaflets were counted and averaged. (B) Difference $\Delta n = n_{\text{upper}} - n_{\text{lower}}$ in the average number of lipids within the square patch around FAM134B-RHD as a function of the leaflet asymmetry $\Delta N = N_{\text{upper}} - N_{\text{lower}}$ for simulations at $\Delta N = -14, 100, 200, 300$. A straight-line fit (orange) gives $\Delta n = -18.1 + 0.114\Delta N$. In a large membrane, we expect $\Delta n = \Delta N$ (dashed grey line) on average, such that the lipid asymmetry of the membrane exactly compensates for the asymmetric footprint of the protein. From the intersection of $\Delta n = \Delta N$ with the linear fit, we find that at equilibrium, FAM134B-RHD displaces 20.4 lipids more from the upper leaflet than from the lower leaflet (red circle).

### Movie Legends

#### Supporting Movie S1

MD simulation trajectory of membrane budding event induced by FAM134B-RHD. A simulation of nine FAM134B-RHD molecules embedded in a membrane with asymmetry $\Delta N = 300$ is shown. The movie covers a time of 2.5 $\mu s$. The same trajectory is analyzed in Figures 1, 3 and 5 of the main text. Phosphate groups are shown in light blue, and lipid tails in white. The proteins are colored in green, orange, yellow and grey, respectively (same coloring as in Figure 5 of the main text). In each frame, the trajectory was centered on the highest phosphate (PO4) bead in $z$-direction, as described in the Methods section.
